# Fibrillation of the *Pseudomonas* Extracellular Functional Amyloid FapC is Controlled by Oligomeric State, Hierarchical Architecture and the C-Terminus

**DOI:** 10.64898/2026.04.28.719052

**Authors:** Chang-Hyeock Byeon, Abdulkadir Tunc, Mustafa Ulasli, Xiangting Tang, Ümit Akbey

## Abstract

Functional amyloids enable bacteria to harness the exceptional stability of the amyloid fold while preventing uncontrolled aggregation. However, the molecular mechanisms that encode this balance between robustness and regulation remain incompletely understood. Here, we define how the *Pseudomonas* functional amyloid FapC, a secreted extracellular functional amyloid in bacterial biofilm, achieves controlled fibril assembly through a modular architecture and a short C-terminal regulatory element. Using controlled purification, modular truncation constructs, and quantitative kinetic assays, we reveal that full-length FapC fibrillation, comprising three irregular layers, is highly sensitive to its initial oligomeric state. Non-monomeric species exert opposing effects: high-molecular-weight assemblies accelerate aggregation via heterogeneous nucleation, while low-molecular-weight species potently inhibit productive assembly. In contrast, isolated single Layer 3, a structurally defined β-solenoid segment, fibrillates rapidly with minimal lag and is largely insensitive to oligomeric heterogeneity, identifying it as a dominant amyloidogenic unit. Yet, despite its minimal architecture, Layer 3 still benefits from the presence of the C-terminal region, which enhances fibrillation rate and yield. Removal of this C-terminal segment delays nucleation, slows elongation, and reduces final fibril formation in both full-length and Layer 3 contexts, highlighting its role as a non-structural regulator of productive folding. Comparative studies show that hierarchical layering imposes kinetic checkpoints and expands regulatory potential, tuning the timing and efficiency of assembly. Together, our results establish that FapC encodes a fibrillation mechanism built on hierarchical architectural design and localized sequence-specific regulation. This suggests that precise control, rather than mere aggregation propensity, maybe the defining hallmark of functional amyloids, distinguishing them from their pathological counterparts.

## 1. Introduction

Amyloid fibrils are among the most stable protein assemblies in nature.^1,2^ Although amyloid formation is classically associated with neurodegenerative disease,^3-5^ it is now well established that many organisms deliberately exploit the amyloid fold for beneficial purposes.^4-11^ In bacteria, functional amyloids in extracellular matrix (ECM) play essential roles in adhesion, biofilm formation, and environmental persistence.^12-14^ These systems raise a central biological question: how can cells harness the extreme stability and self-assembly propensity of amyloids while preventing uncontrolled aggregation that would compromise cellular function?

A defining feature of functional amyloids is that their assembly is precisely regulated in space, time, and efficiency. In contrast to pathological amyloids,^3^ which often arise from loss of proteostasis, functional amyloids are produced through genetically encoded intrinsic/extrinsic pathways that coordinate expression, secretion, and controlled extracellular assembly.^15-26^ Despite recent structural advances revealing the architectures of mature functional amyloid fibrils, the molecular principles that govern their controlled assembly remain incompletely understood. In particular, the determinants that regulate the transition from a disordered monomeric state to a highly ordered cross-β solenoid architecture are largely unresolved.

The functional amyloids in *Pseudomonas* species (Fap) provides a powerful model to address these questions. FapC, the major amyloid subunit of the biofilm matrix, is an intrinsically disordered protein (IDP) in its monomeric state that undergoes a disorder-to-order transition upon fibril formation.^27,28^ Secreted into the ECM, FapC assembles into amyloid fibrils that confer biofilm integrity, mechanical stability and thus resilience to antibiotics. Because FapC-mediated biofilm formation contributes to bacterial persistence and antimicrobial tolerance, understanding its aggregation mechanism is directly relevant to strategies aimed at disrupting biofilm-associated chronic infections.^29-32^

FapC is encoded within the Fap operon composed of six proteins, FapA-F,^33^ including accessory proteins involved in secretion, proteolytic processing, transport and nucleation, underscoring that Fap amyloid assembly is tightly regulated at multiple levels. We have recently shown by high-resolution structural studies that mature full-length FapC (25-250, FL) fibrils adopt an ordered cross-β-solenoid architecture (PDB: 9NQD & EMDB: 49649).^27^ This β-solenoid fold is composed of three stacked β-rich layers derived from regions spanning approximately residues 38-99 (Layer 1), 100-164 (Layer 2), and 165-236 (Layer 3, L3). These layers are flanked by flexible N- and C-terminal segments that do not form part of the rigid amyloid core but forming a fuzzy coat. Notably, the short N/C-terminal regions lie outside the ordered cross-β core and remains conformationally dynamic, may have a potential regulatory function rather than structural role in fibrillation.^27^

While the structural organization of mature FapC fibrils is now established, structure alone does not explain how assembly is kinetically controlled. Our prior work has suggested that functional amyloid formation may proceed through hierarchical or stepwise pathways and that non-core regions (disordered and/or fuzzy-coat) can influence nucleation and elongation.^27,34^ However, direct experimental separation of how modular layer architecture, oligomeric starting states, and sequence elements cooperate to regulate fibrillation kinetics has been lacking. In particular, the contribution of the C-terminal segment to assembly remains ambiguous, and although it does not form part of the fibril core, it may influence fibril formation and so biofilm integrity.

Building directly on our FapC fibril structure, we systematically analyze its fibrillation mechanism by separating three variables that are often mixed in amyloid studies: heterogenic starting state, modular layer architecture, and C-terminal sequence context. Using controlled purification strategies, size-exclusion chromatography (SEC), and a comprehensive set of truncated and modular FapC constructs (FL vs. L3), we directly compare fibrillation kinetics, lag behavior, concentration dependence, and final fibril yield under defined assembly conditions.

By isolating individual layers and selectively removing the C-terminal region, we demonstrate that FapC fibrillation follows a hierarchical and regulated assembly mechanism. A single layer, L3, functions as a dominant amyloidogenic module capable of rapid, robust fibrillation, whereas FL FapC exhibits kinetic checkpoints that impose a lag phase and sensitivity to oligomeric heterogeneity. We further show that non-monomeric species exert opposing effects on fibrillation, acting either as heterogeneous nucleation sites or as inhibitors of productive assembly depending on their size and architectural context. Importantly, the C-terminal region is not required for amyloid formation itself but is essential for achieving efficient, high-yield fibrillation, suggesting it as a regulatory element rather than a structural component of the amyloid core. Together, these findings establish a mechanistic framework in which functional amyloid assembly is encoded through modular architecture and sequence-specific regulation, enabling robust fibril formation while maintaining kinetic control. This work clarifies how bacteria engineer amyloid materials that are both structurally stable and precisely regulated, and suggests regulation, not aggregation propensity, as the defining feature distinguishing functional from pathological amyloids.

## 2. Methods

### FapC Constructs

The FapC 25-250 (FL, without the signal sequence comprising 1-24) preparation has been described previously in detail.^27,35^ FapC FL was previously cloned into a pET28a vector system with a C-terminus His-tag motif. All other clones were generated using a modified pET28a vector, with improvements for expression elements and a TEV cleavage site instead of thrombin, see full protein sequences in **Fig. SI1**. The T7pcons and TIR-2 modified sequences in the T7 promoter and translation initiation region were used to enhance protein expression.^36^ FapC 25-236 (FL ΔC) was cloned into the modified pET28a vector with a C-terminal His-tag. FapC 165-250 (L3) and 165-236 (L3 ΔC) were cloned into the same vector with an N-terminal His6-tag and TEV cleavage site, with no changes at the C-terminus.

### FapC Expression and Purification

The preparation of FL FapC has been described previously.^27,35^ Briefly, recombinant FapC constructs were expressed in *E. coli* BL21(DE3) cells. Unless otherwise noted, proteins were expressed unlabeled for biochemical and kinetic assays. For isotopically labeled samples used in NMR experiments, cells were grown in minimal medium with ^15^N-ammonium chloride (1 g L^−1^) as the sole nitrogen source; unlabeled samples were grown in LB medium. Overnight LB cultures containing kanamycin were inoculated from glycerol stocks and grown at 37 °C with shaking. Cultures were harvested and resuspended in fresh medium and grown until OD600 reached 0.8-1.0. Protein expression was induced with 1 mM IPTG and continued for 2 h at 37 °C. Cells were harvested by centrifugation (6000 × g, 20 min, 4 °C) and resuspended in denaturing lysis buffer (50 mM Tris pH 8.0, 8 M guanidinium chloride). Cells were lysed by two rounds of sonication (4 min total; 60% power, 50% duty, 5 s cycles) on ice. Insoluble debris was removed by centrifugation (6000 × g, 20 min, 4 °C) to yield clarified lysate. His-tagged FapC was purified under denaturing conditions using Ni-NTA resin (HisPur, Thermo Fisher). Lysates were incubated with resin overnight at 4 °C with gentle mixing. After washing with lysis buffer, proteins were eluted stepwise using 25 mL of lysis buffer containing 30, 60, 120, 300, or 500 mM imidazole. Elution fractions were analyzed individually and pooled as indicated. Proteins were concentrated using 3 kDa MW cut-off Amicon stir cells (Millipore-Sigma). Unless otherwise noted, references to Ni-NTA-purified protein indicate the 300-500 mM elution.

### Size-Exclusion Chromatography (SEC) Separation of Monomer and Non-monomer Species

SEC was used to fractionate FapC samples into defined oligomeric populations before fibrillation assays. Combined Ni-NTA elutions were loaded onto a pre-equilibrated HiLoad 16/600 Superdex 200 column (Cytiva) in lysis buffer containing 8 M guanidinium hydrochloride to maintain denatured conditions. Protein was loaded at concentrations optimized for resolution between monomeric and oligomeric species. Elution was monitored at 280 nm, and fractions were grouped into four pools: (i) high-molecular weight (HMW) near void volume, (ii) medium-molecular weight (MMW), (iii) predominantly monomeric, and (iv) low-molecular weight (LMW) post-monomer peak. These definitions were consistently applied across constructs and replicates. Pools were concentrated to 100-500 μM, flash-frozen, and stored at -80 °C.

### Mass Spectrometry (MS) data acquisition and processing

ESI LC-MS measurements were performed at 1 μM protein concentration on a Bruker Q-TOF instrument, using a reverse phase AdvanceBio peptide guard column (Agilent Technology) where mobile phases A and B comprised 5% and 80% acetonitrile with 0.01% formic acid, respectively. Bruker Compass Software was used to process the LC-MS spectra, and the MS data were processed using maximum-entropy-based deconvolution to obtain the M+ ion mass for each sample. The instrument was calibrated using the ESI Low Tuning mix I (Agilent Technologies) to 1.0 ppm mass % difference before each use.

### Aggregation Kinetics Assays

FapC samples stored in 8 M guanidinium hydrochloride were desalted into fibrillation buffer (20 mM sodium phosphate pH 7.8, 0.02% sodium azide) using PD MiniTrap G-10 columns (Cytiva). Protein concentration was determined by UV absorbance at 280 nm using NanoDrop (Thermo Scientific) and the following extinction coefficients: FapC 25-250: 10.1 mM^−1^ cm^−1^, FapC 25-236: 15.47 mM^−1^ cm^−1^, FapC 165-250: 9.97 mM^−1^ cm^−1^, FapC 165-236: 9.97 mM^−1^ cm^−1^. These coefficients were used to estimate total protein concentrations, with the caveat that cleaved or oligomeric species (except in monomeric fractions) reducing effective monomer concentration. FapC was diluted to 10-100 μM in fibrillation buffer, with 200 μM thioflavin T (ThT) added. 50 μL aliquots were loaded into black, clear-bottom 96-well plates (Corning 3881, Sigma-Aldrich). Fluorescence was monitored in a Tecan Spark reader (excitation: 442 nm; emission: 482 nm) every 10 min for 65 h at 37 °C without shaking. All assays were run in triplicate. Background ThT fluorescence (with buffer only) was subtracted. To assess fibrillation by SDS-PAGE, samples were collected before (PRE) and after (POST) ThT incubation. PRE samples were mixed with SDS-loading buffer and denatured at 50 °C before storage at 4 °C. POST samples were prepared similarly. Both sets were run under non-reducing conditions on 12% SDS-PAGE gels. Monomer band intensity was quantified in ImageJ and expressed as a percentage of starting monomer in PRE lanes.

### Solution-State NMR Spectroscopy

Uniformly ^15^N-labeled FapC monomer was used for NMR. Spectra were acquired on a Bruker Avance 700 MHz spectrometer with 5-mm TCI cryogenic probes. Samples contained 50 μM protein in 20 mM sodium phosphate pH 7.8, 0.02% sodium azide, 1 mM deuterated dextran sulfate sodium (DSS), and 10% D_2_O in 420 μL volumes. Measurements were collected at 274 K. Assignments were adapted from previous work.^35^ Spectra were processed in TopSpin 3 and assigned in CCPNmr Analysis 3. Proton shifts were referenced to DSS (0 ppm), and ^15^N shifts were indirectly referenced using ^1^H frequency.^37^ Chemical shift perturbations (CSPs) were calculated using Δδ = ((5*Δδ^1H^)^2^ + (Δδ^15N^)^2^)^1/2^ and grouped as: no CSP (<0.035 ppm), moderate CSP (0.035– 0.07 ppm), or strong CSP (>0.07 ppm).

### Negative-Stain Electron Microscopy (EM)

Negative-stain EM was used to examine fibril morphology and abundance. Copper 400-mesh grids with continuous carbon film were glow discharged for 90 s. 3 μL of sample was applied, incubated 10 s, and blotted with filter paper. Grids were stained with 2% (w/v) uranyl acetate for 10 s, then blotted and air-dried. Micrographs were acquired using a Tecnai TF20 transmission electron microscope (Thermo Fisher) at 200 kV with a TVIPS XF416 CMOS camera. No image post-processing was applied. Fibril widths were measured using ImageJ from the negative-stain EM micrographs. All the EM micrographs shown have 200 nm scale bar.

## 3. Results

### FapC functional amyloid structure and modular β-solenoid layer organization

The cryo-EM structure of fibrillar FapC we recently presented, **Fig. 1A-C**, reveals a three-layered cross-β-solenoid amyloid fibril fold,^27^ which provides structural basis for how an intrinsically disordered protein transitions into a stable functional amyloid. Similar to extracellular functional amyloid CsgA, and intracellular pathologic amyloids HELLF and HetS, FapC also adopts a β-solenoid fold.^38-40^ Each FL FapC monomer unit in the amyloid fold is constituted from three β-sheet-rich layers, Layer 1 (residues 38-99), Layer 2 (100-164), and Layer 3 (165-236), which together form a continuous β-solenoidal amyloid structure, **Fig. 1B,D**. The FapC fibril layers include both conserved imperfect repeats (R1-R3) and intervening non-repeat loops (L1, L2),^34^ suggesting that both motif types are critical for structural organization. Biochemical and mutagenesis studies, including truncation of loops or individual repeat units, support this conclusion, that the consensus and non-consensus regions are essential for fibril.^34,41^ The layers are connected by flexible turns of varying lengths and flanked by highly disordered N- and C-terminal tails (residues 25-37 and 237-250). These flexible and disordered regions remain unresolved in cryo-EM density but we previously studied these by solution NMR and all-atom molecular dynamics (MD) simulations.^27^ The FapC fibril forms a single protofilament with an axial rise of ∼14.6 Å per monomer (∼4.85 Å cross-β layer distance and three layers), **Fig. 1B**. Each layer contains five β-strands forming a rigid S-shaped β-sheets core, flanked by two disordered β-sheets. Cryo-EM density reveals alternating rigid and flexible regions, resulting in a fibril with structurally heterogeneous regions. This observation is supported by solid-state NMR spectra, which distinguish pools of rigid and flexible residues.^27^ Our prior NMR assignments of monomeric FapC confirmed its intrinsically disordered character and revealed local secondary structure propensities that match with secondary structure elements in the AlphaFold2 (AF2) model (AF-C4IN70-F1), **Fig. 1B, SI1**. The AF2 model adopts an irregular 2-1-3 layer topology, although ∼50% of its structure is predicted with low confidence supported by our cryo-EM structure using CryoSPARC.^27^ In contrast, the AlphaFold3 (AF3) model (AF-P0DXF5-F1) proposes a monomeric fold with a regular 1-2-3 layer order, **Fig. SI2**.^42^ This divergence likely arises from increased intramolecular constraints emphasized in AF3. These discrepancies point out that that predictive models prioritize different structural features depending on training objectives and need for structural validation.

**Figure 1.**
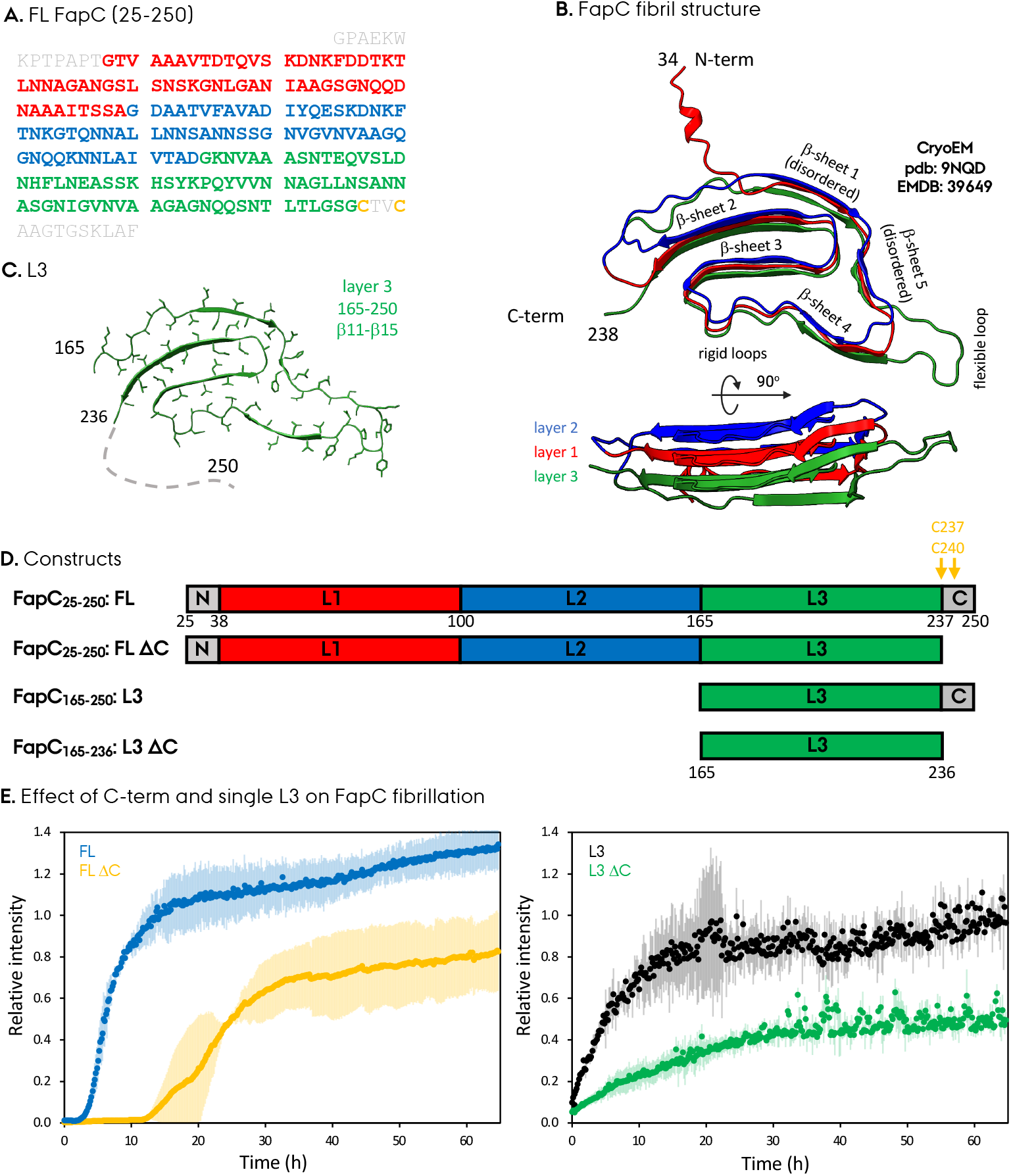
Structural framework and modular organization of FapC fibrils. (A) Schematic of full-length (FL) FapC showing three β-solenoid layers (Layers 1-3, L1-3), linker regions, and disordered C-terminal segment. (B) Cryo-EM model of mature fibril structure revealing cross-β architecture with a rigid amyloid core and flexible outer regions. (C) Diagram of isolated L3 region from the EM model; dashed C-terminal region indicates unresolved density. (D) Construct overview: FL and L3, with and without the C-terminal segment. (E) Deletion of the C-terminus reduces fibrillation efficiency for FL and L3, shown by ThT assay at 40 μM using pure monomeric samples.

Collectively, our cryo-EM data, AF2 model and all-atom MD simulations support an interleaved irregular 2-1-3 three-layer β-solenoid fold for FapC, **Fig. 1B, SI2**. In the next sections, we describe how fibrillation kinetics are modulated by layer composition, sample purity, oligomeric state, and the C-terminal region using tailored constructs. The distinct ThT profiles shown in **Fig. 1E** for all four FapC constructs suggest the functional role of C-terminal and the layers on fibrillation.

### Purity and oligomeric state influence fibrillation efficiency of FapC

The initial purity of monomeric proteins has a major effect on fibrillation kinetics, as subtle compositional differences can alter aggregation. Previous studies, including our own, have predominantly relied on Ni-NTA affinity purification to isolate FL FapC (residues 25-250, **Fig. 1A,B**). While this method yields relatively pure protein (>95%), SDS-PAGE analysis of the Ni-NTA elution consistently revealed the presence of non-monomer species, **Fig. 2A,B**. Alongside the expected monomeric band at 23.7 kD, we observed additional high-molecular-weight (HMW) species (composed of FapC oligomers), medium-molecular-weight (MMW) species (such as dimers), and low-molecular-weight (LMW) species (cleavage products), **Fig. 2B**. Importantly, the relative abundance of these species varied across the Ni-NTA imidazole elution gradient. This indicates that the conventional Ni-NTA purification protocol does not yield a biophysically homogeneous FapC sample. To improve sample definition in terms of purity as well as oligomeric status, we implemented a secondary purification step using SEC, **Fig. 2C**. This approach yielded a highly pure monomeric fraction as also seen in the MS of the monomeric SEC fraction, **Fig. 2D**, as well as fractions containing monomer mixed with well-defined LMW, MMW, and HMW species. By isolating these populations, we established a clear framework for evaluating how specific oligomeric states influence FapC fibrillation, discussed in the following sections.

**Figure 2.**
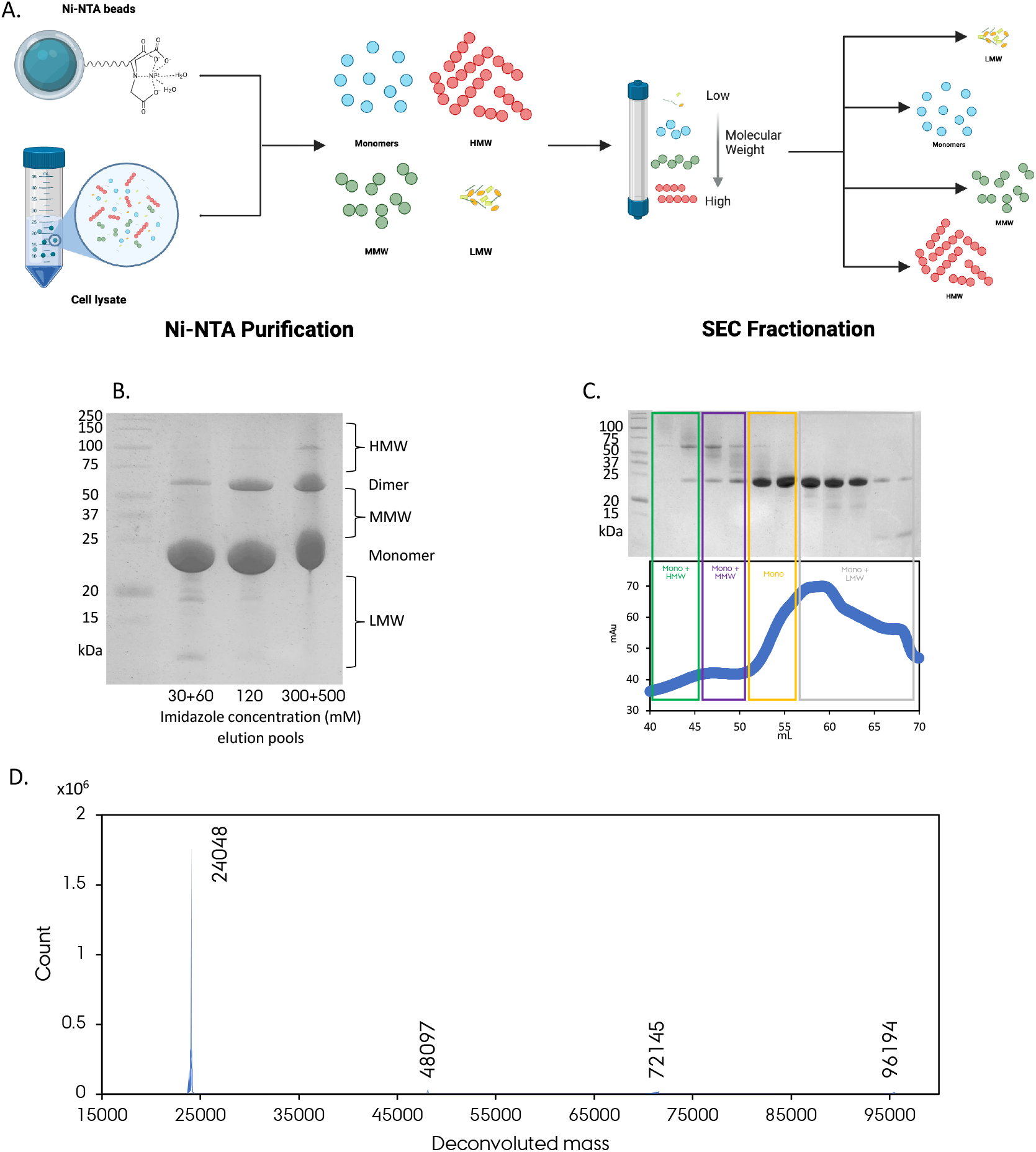
Heterogeneity in affinity-purified FL FapC and isolation of distinct oligomeric states by SEC. (A) Purification schematic we used for our samples. (B) Ni-NTA purification of FL FapC (25–250) yields mixed oligomeric species. (C) SEC separates these into defined monomeric and non-monomeric pools, enabling controlled assessment of their impact on fibrillation behavior. (D) Mass spectrometry result of a ^15^N FapC FL from SEC Mono+MMW fraction shows ∼95% ^15^N labeling with most abundant monomer as well as observable dimer, trimer and tetramer species.

### Non-monomeric species modulate full-length FapC fibrillation

ThT assays performed on Ni-NTA elution fractions revealed distinct differences in fibrillation kinetics, **Fig. 3A-C**. Lower-purity 30-60 mM and 120 mM imidazole fractions (samples #2 and #3) displayed extended lag phases and significantly reduced ThT fluorescence, consistent with inefficient fibrillation due to impurities. In contrast, the higher-purity 300-500 mM imidazole fraction (sample #1) showed rapid assembly with a fibrillation lag time (T_lag_) of 3 hours (h) and fibrillation half time (T_50_) of 8 h, **Fig. 3A**. SDS-PAGE gels supported these observations: only ∼10% and ∼20% monomer depletion occurred in samples #2 and #3, while sample #1 showed ∼90% monomer reduction, **Fig. 3B**. EM images corroborated this trend, and sample #1 showed abundant fibrils with minimal amorphous aggregates, while samples #2 and #3 displayed predominantly amorphous material, **Fig. 3C**. We observed an inverse correlation between amorphous aggregates observed in EM and ThT fluorescence. Moreover, the amount of amorphous aggregates correlated with sample heterogeneity.

**Figure 3.**
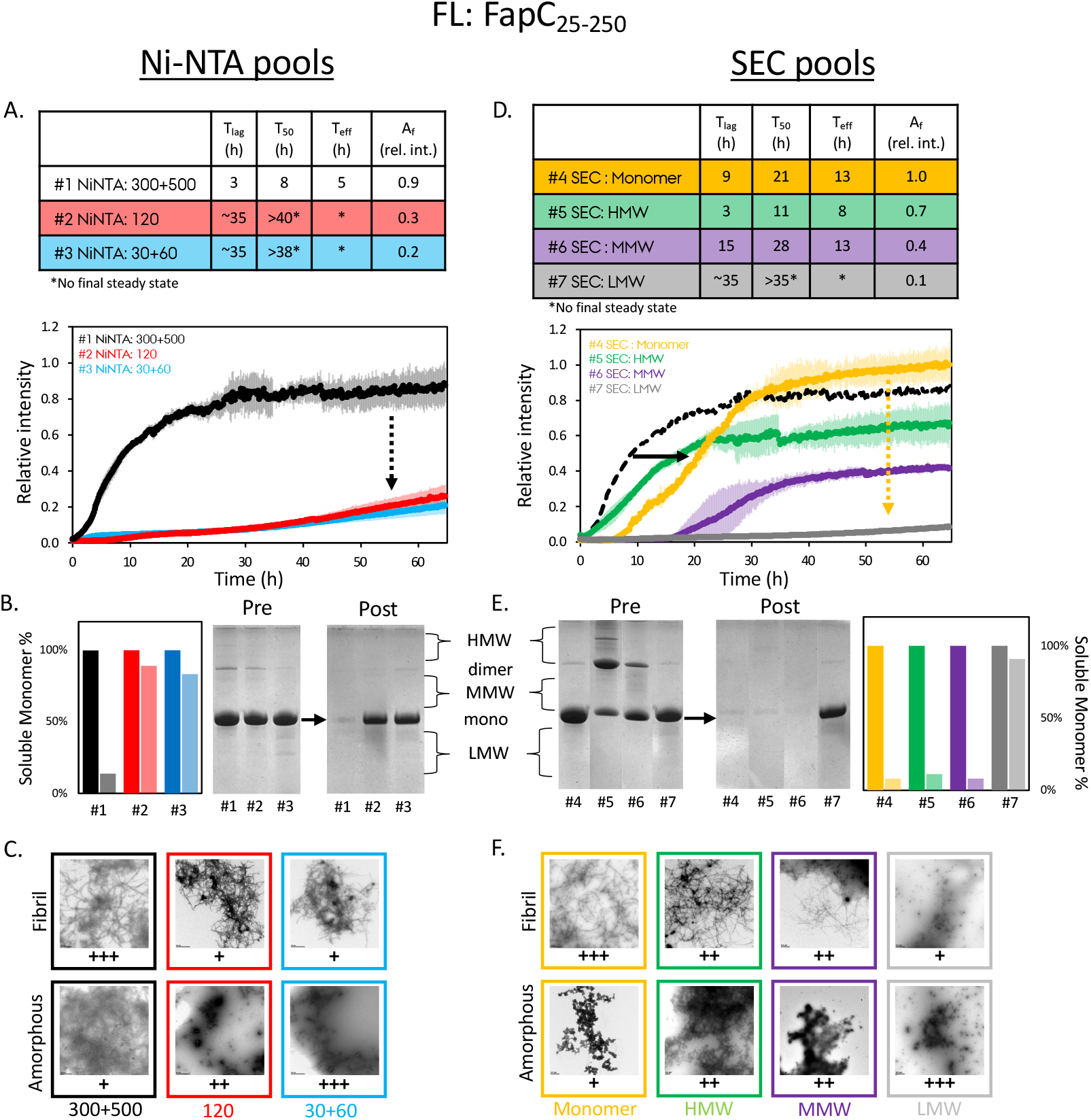
Distinct non-monomeric species modulate FL FapC fibrillation. Ni-NTA and SEC fractions (samples #1–7) were characterized by ThT fluorescence, SDS-PAGE, and EM. All samples were measured 40 μM based on UV280 quantification of desalted sample. The ThT fluorescence was normalized relative to SEC monomer (sample #4). (A, D) ThT traces show varying fibrillation kinetics depending on oligomeric content for Ni-NTA pools (samples #1-3) and SEC pools (samples #4-7). ThT samples that did not reach saturation were labeled with an asterisk and values are all approximations. In D, The ThT of sample #1 that is shown in A is additionally added for comparison. (B, E) SDS-PAGE gels of PRE and POST samples illustrate monomer loss. (C, F) EM images show fibril and amorphous aggregate abundance; “+” indicates qualitative enrichment. ThT intensity and SDS-PAGE monomer depletion correlate with fibril presence. The arrows indicate the effect of non-monomeric FapC species to the ThT profiles.

To identify the role of specific non-monomeric species, we analyzed SEC-separated pools from the same Ni-NTA-purified FapC. All four SEC pools contained monomer, **Fig. 3D-F**, allowing us to assess the impact of co-eluting species. Three of the four SEC pools (samples #4-6) achieved near-complete fibrillation with >90% monomer depletion by SDS-PAGE, **Fig. 3E**. Oligomeric species in these fractions also disappeared post-incubation, suggesting incorporation into fibrils or amorphous aggregates, **Fig. 3E**.

The monomer-only pool (sample #4) showed a longer T_lag_:9 h and T_50_:21 h, while reaching a final ThT signal comparable to NiNTA:300+500 sample #1. Given no detectable oligomers, we regard sample #4 as the baseline intrinsic FapC fibrillation absent external modulators, as a reference. The difference between sample #1 and #4 highlights that the initial oligomeric state influences nucleation, lag phase, and elongation. Oligomers present in Ni-NTA purified sample appear to facilitate seeding and substantially reduce the lag phase. This resembles previously observed vessel surface effects, where removal of surface contact via microdroplets extended lag and elongation phases.^25^ These findings suggest that FapC fibrillation may benefit from both surface-mediated nucleation and oligomer-driven seeding, and may suggest the need for both to have the reduced lag phase. Supporting this, the HMW pool (sample #5) exhibited similar kinetics to sample #1 (T_lag_:3 h, T_50_:11 h) while achieving ∼75% of the final ThT signal. The MMW pool (sample #6) showed intermediate kinetics between monomer and HMW pools. EM confirmed these trends, and sample #4 showed dense fibrils with minimal aggregates, but samples #5 and #6 displayed mixed morphologies, **Fig. 3F**. Remarkably, the LMW pool (sample #7) was highly deficient in fibrillation. Only ∼10% monomer reduction was observed by SDS-PAGE, with ThT signal at just 10% of sample #4 and delayed kinetics (T_lag_ ∼35 h). This mirrors low-purity Ni-NTA fractions (samples #2 and #3). The presence of LMW species appears to inhibit FL FapC fibrillation, likely by sequestering monomer or promoting off-pathway aggregation. EM confirmed sparse fibrils and abundant amorphous aggregates.

These results demonstrate that the identity, not just the presence, of coexisting species profoundly impacts FL FapC fibrillation. HMW oligomers likely promote fibrillation by bypassing nucleation barriers, while LMW species inhibit productive assembly. EM morphologies track closely with ThT profiles, reinforcing that oligomeric heterogeneity can qualitatively reshape FapC assembly. This underlines the importance of precise sample characterization when studying functional amyloids.

### Minimalistic Layer 3 fibrillates efficiently regardless of oligomeric state

To determine the fibrillation efficiency of the minimal single-layer construct L3, we examined its behavior across Ni-NTA and SEC purification fractions and compared it with FL FapC. L3 was selected based on our previous findings identifying it as the most aggregation-prone region of FapC.^27^ Even in the monomeric IDP state, L3 shows strong intrinsic β-sheet propensity as monomer and contributes significantly to the fibril core.^35^ We also demonstrated by MD simulations that L3 undergoes rapid conformational transitions during early assembly, often preceding other regions in adopting β-solenoid structure.^27^ L3 acts as a nucleation point to initiate fibrillation by stabilizing early folding intermediates, which was also seen on other amyloids, e.g. insulin, where specific regions act as “hotspots’ for nucleation.^43^ This made L3 an ideal minimal system for assessing core amyloid-forming capacity without hierarchical regulation by other layers. L3 was purified by Ni-NTA and SEC, enabling comparisons under matched conditions, **Fig. 4A**.

**Figure 4.**
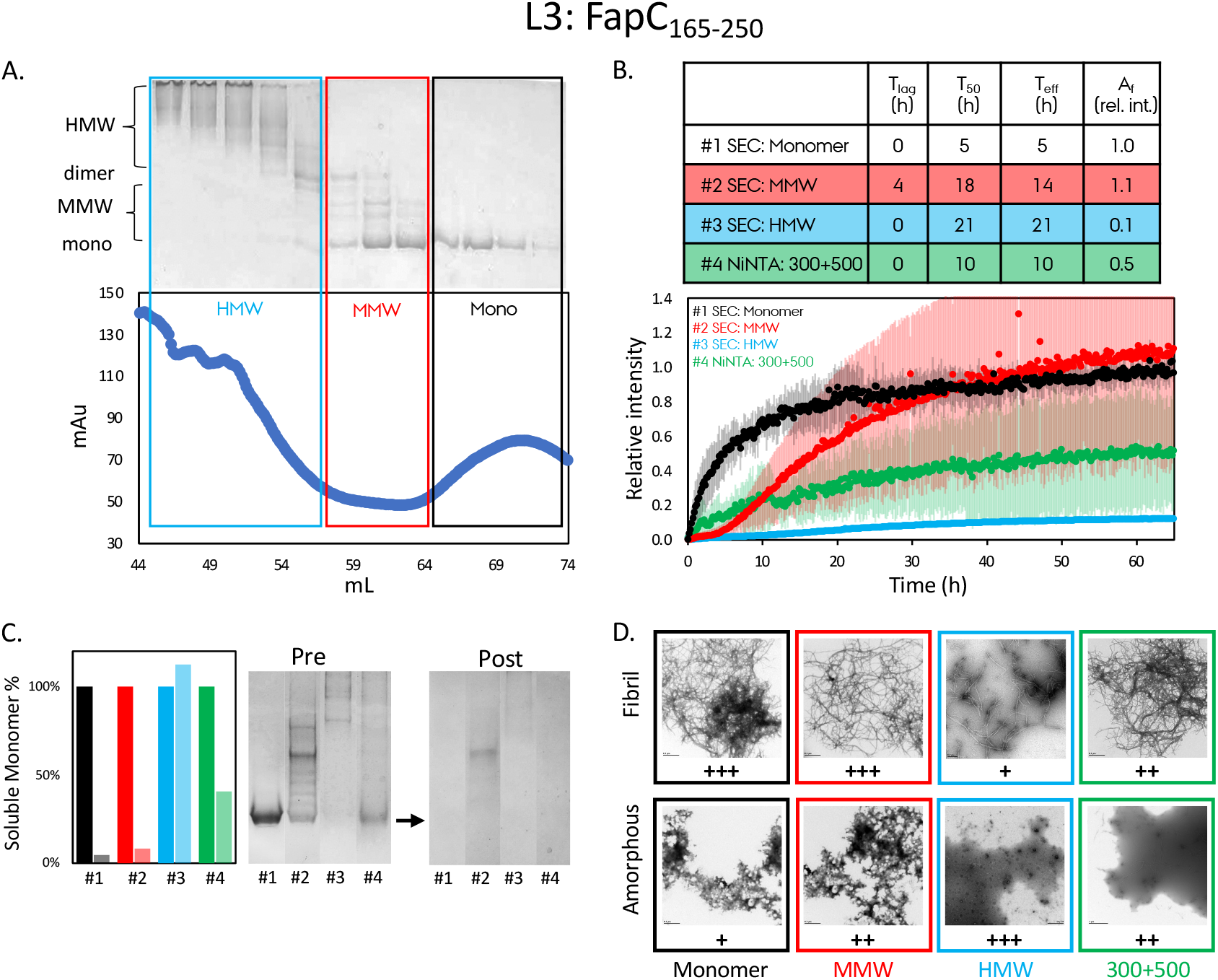
L3 fibrillation is robust and minimally affected by oligomeric heterogeneity. L3 samples purified by Ni-NTA and SEC were analyzed by ThT, SDS-PAGE, and EM. All samples were measured at 50 μM based on UV280 quantification of desalted sample. The ThT fluorescence was normalized relative to SEC monomer (sample #1). (A) SEC and SDS-PAGE of pooled L3 fractions: monomer, HMW, and MMW samples. (B) ThT profiles correlate with monomer content showing rapid fibrillation across samples that is mildly affected by non-monomer species. (C) SDS-PAGE images of PRE and POST confirms monomer depletion due to fibrillation and consistent with the ThT assay results. Sample #3 PRE monomer quantification by ImageJ was very low and resulted in a relatively large error. (D) EM reveals fibrils and amorphous aggregates across all samples, and fibril content mirrors ThT and SDS-PAGE results.

Ni-NTA-purified L3 (300+500 mM elution; sample #4) fibrillated rapidly, with no detectable lag phase and T_50_:10 h. SDS-PAGE showed a significant (∼60%) decrease in monomer signal, **Fig. 4C**, and EM revealed abundant fibrils and moderate amorphous aggregates, **Fig. 4B-D**. The SEC-purified monomer-only L3 pool (sample #1) also fibrillated rapidly, with no lag and a shorter T_50_:5 h. This sample showed faster fibrillation than the L3 Ni-NTA pool, FL Ni-NTA pool, and FL monomer sample, **Fig. 4C**. EM confirmed fibrils with some aggregates, **Fig. 4D**. The L3 MMW pool (sample #2) exhibited T_lag_:4 h and prolonged T_50_:18 h, compared to monomer-only L3. This trend mirrored FL MMW vs. FL monomer samples, **Fig. 3E**. In both, SDS-PAGE showed ∼90% monomer reduction along with disappearance of oligomeric species. EM confirmed fibrils with moderate amorphous aggregates, **Fig. 4D**. The oligomer-only HMW pool (sample #3) showed no monomer band on SDS-PAGE prior to ThT, low ThT fluorescence (∼10%), and sparse fibrils with dominant aggregates by EM, **Fig. 4D**. Notably, unlike FL FapC, L3 samples lacked detectable LMW species.

In contrast to FL, L3 fibrillation was largely insensitive to coexisting oligomers. This insensitivity supports the idea that L3 can nucleate and elongate efficiently without external facilitation, and that nucleation is not rate-limiting in this minimal context. The only exception was the MMW pool, which delayed elongation but reached a comparable ThT endpoint. These observations support a hierarchical model in which minimal modules like L3 fibrillate autonomously, while higher-order architecture in FL adds kinetic regulation. These results demonstrate that L3 is an intrinsically efficient amyloid-forming module with fibrillation governed mainly by monomer availability rather than structural regulation. Unlike FL, which requires proper folding of three β-solenoid layers, L3 assembles rapidly and robustly even in the presence of oligomeric heterogeneity.

### Disordered C-terminus promotes FapC amyloid assembly

To investigate the role of the conformationally flexible C-terminus in fibrillation kinetics and efficiency, we deleted residues 237-250 (CTVCAAGTGSKLAF) from both FL and L3 constructs, generating FL ΔC (25-236) and L3 ΔC (165-236) variants, **Fig. 1E, 5**. This region lies outside the β-solenoid core but may still influence assembly by being part of the fuzzy-coat, and contains two cysteine residues that may influence fibrillation, **Fig. 1A-D**. ^27,44^ C-term deletion constructs were purified using Ni-NTA and SEC protocols, and their fibrillation was compared using ThT assays on SEC-purified monomer-only fractions, **Fig. 1E, 5**. Across both backgrounds, C-terminal deletion consistently reduced final ThT fluorescence, **Fig. 1E**, increased T_50_, and in FL prolonged the lag phase, supporting a positive contribution of the C-terminal segment to efficient fibrilization, **Fig. 5**. Flexible C-termini affecting fibrillation was observed for pathologic amyloids, e.g. for αSyn, aβ42, where the C-terminal residues are important for fibril formation, whereas similar C-termini inhibits Tau fibrillation.^45-47^

**Figure 5.**
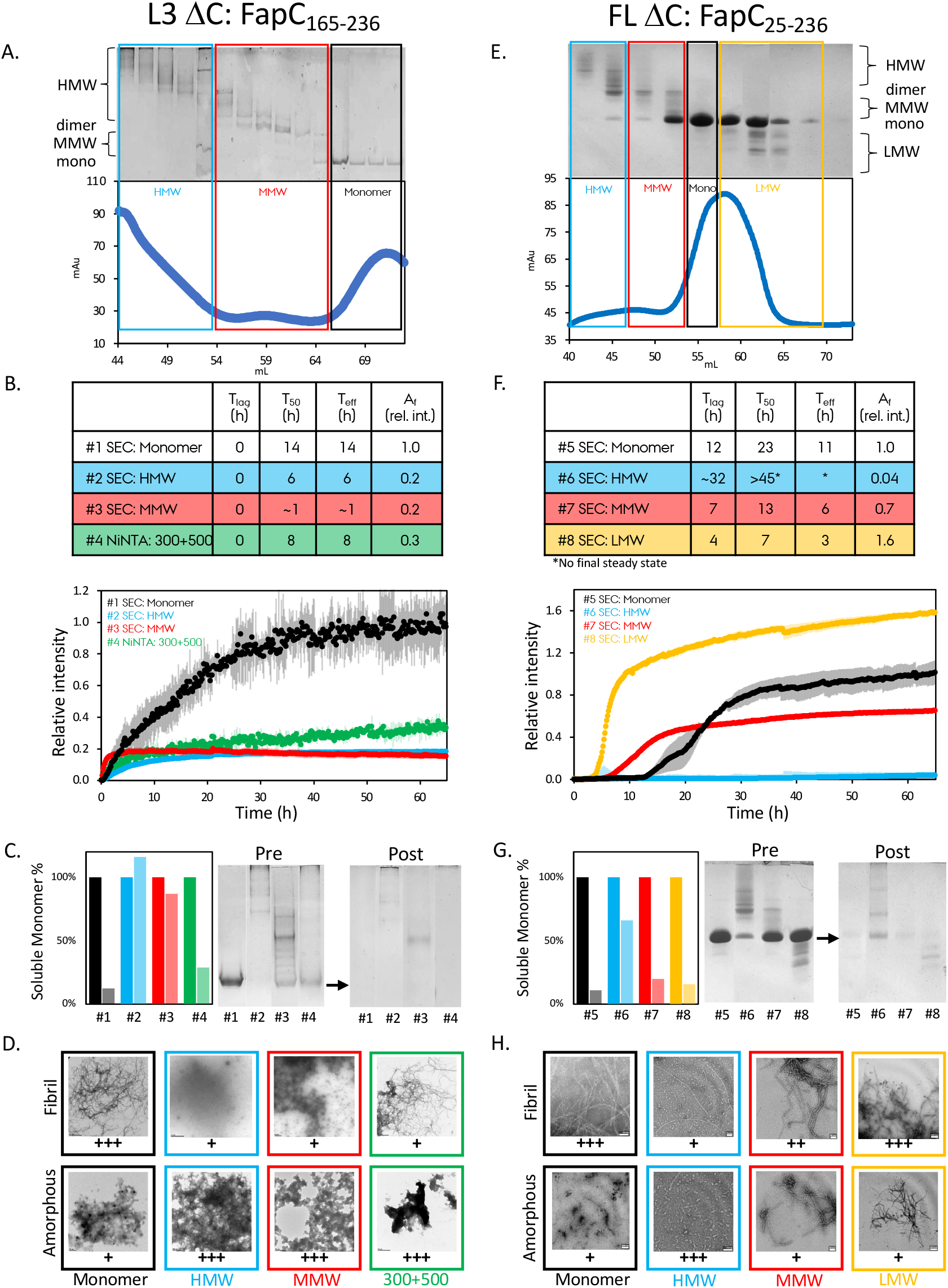
Removal of the C-terminus alters fibrillation efficiency and sensitivity to assembly state. Deletion of the C-terminal region in FL and L3 affects fibrillation kinetics, oligomer sensitivity, and fibril morphology. SEC-purified samples were analyzed by ThT, SDS-PAGE, and EM. Samples were measured at 40 μM (L3 ΔC) and 50 μM (FL ΔC), respectively, based on UV280 quantification of desalted samples. The ThT fluorescence was normalized relative to their respective SEC monomer (sample #1 and #5). (A, E) SEC profiles and the corresponding SDS-PAGE gels for L3 ΔC and FL ΔC. Samples were separated to monomer, HMW, MMW and LMW fractions. (B, F) The ThT profiles for different SEC fractions for L3 ΔC and FL ΔC. ThT samples that did not reach saturation were labeled with an asterisk and values are all approximations. (C, G) SDS-PAGE images of PRE and POST confirms monomer depletion due to fibrillation and consistent with the ThT assay results. Sample #2 PRE monomer quantification by ImageJ was very low and resulted in a relatively large error. (D, H) EM reveals fibrils and amorphous aggregates across all samples, and fibril content mirrors ThT and SDS-PAGE results.

In the L3 ΔC monomer fraction (sample #1), fibrils formed with ∼90% monomer depletion by SDS-PAGE and abundant fibrils in EM, **Fig. 5A-D**. Despite the absence of a lag phase, T_50_ increased from 5 h in L3 to 14 h in L3 ΔC, **Fig. 4B, 5B**, indicating that the C-terminus promotes elongation. The Ni-NTA 300-500 mM elution (sample #4) also lacked a lag phase but showed a reduced final ThT signal, correlates with low initial monomer amount. MMW and HMW fractions exhibited low ThT fluorescence, minimal monomer loss, and sparse fibrils on EM, likely forming off-pathway aggregates with limited ThT binding, **Fig. 5C, D**.

In FL ΔC, a similar trend was observed, **Fig. 5E-H**. MMW and LMW pools retained fibrillation competence but with slower kinetics than FL, **Fig. 5F-H**. The FL ΔC monomer (sample #5) had a T_lag_:12 h and T_50_:23 h, both longer than in FL, **Fig. 5F**. Interestingly, the MMW pool (sample #7) had shorter T_lag_:7 h and T_50_:13 h than its FL counterpart, **Fig. 3D**, suggesting partial compensation by oligomeric species. Surprisingly, the LMW pool (sample #8) showed enhanced fibrillation, 60% higher ThT signal and faster kinetics (T_lag_:4 h, T_50_:7 h), in contrast to the inhibitory behavior of LMW species in FL, **Fig. 3D, 5F**. This implies that the C-terminal segment may interact differently with LMW, modulating their effect on fibrillation. The HMW pool of FL ΔC (sample #6) showed negligible ThT and few fibrils by EM, correlates with low monomer amount, mirroring the HMW behavior in L3 and L3 ΔC.

Together, these findings demonstrate that the disordered C-terminal region is important in FapC and enhances efficient fibrillation in both FL and L3 contexts. Its influence is context-dependent: deletion reduces fibrillation efficiency in monomer-only samples and alters how oligomeric species affect assembly. These results suggest that the C-terminal segment contributes to conformational selection and alignment of subunits during early nucleation and elongation.

### Concentration-dependent kinetics reveal non-redundant contributions of hierarchical architecture and the C-terminus

To further understand how FapC assembly responds to physiological variation, we analyzed fibrillation kinetics across a range of protein concentrations for all four constructs (FL, FL ΔC, L3, L3 ΔC), using monomer-only samples purified by SEC. Fibrillization kinetics are highly dependent on monomer concentration, a phenomenon well-documented in amyloid research.^48^ A comparison of all FapC constructs at 40 μM revealed key distinctions, **Fig. 6A**. FL showed a short T_lag_:4 h and T_50_:8 h, while FL ΔC had a significantly longer T_lag_:12 h and T_50_:23 h. When expressed as effective fibrillization time (T_eff_=T_50_-T_lag_), these were 4 and 11 h, respectively, **Fig. 6F,SI3**. L3 and L3 ΔC showed no lag phase, but T_eff_ differed markedly: 6 h for L3 and 14 h for L3 ΔC, highlighting the C-terminus’s contribution to fibrillation efficiency. Final ThT fluorescence at 40 μM was ∼60% higher for FL versus FL ΔC and ∼80% higher for L3 versus L3 ΔC, **Fig. 6A**. Notably, average fibril widths showed variations across constructs, 8.2 nm (FL), 7.4 nm (FL ΔC), 10.2 nm (L3), and 8.9 nm (L3 ΔC), with a standard deviation of 1.7-2.8 nm. The fibril widths for the constructs with C-terminus are ∼1 nm wider compared to the ΔC versions, indicating that C-terminus may have an effect on the fibril width, **Fig. 6B-E, SI4**. Surprisingly, the L3 fibril widths with/out C-terminus are ∼1.5-2 nm wider compared to the FL counterparts. These data confirm that L3 ΔC retains the minimal amyloid-forming unit, while the C-terminus enhances fibrillation speed and yield. Additionally, the hierarchical triple-layer structure in FL slows nucleation, introducing a lag phase that modulates fibril onset.

**Figure 6.**
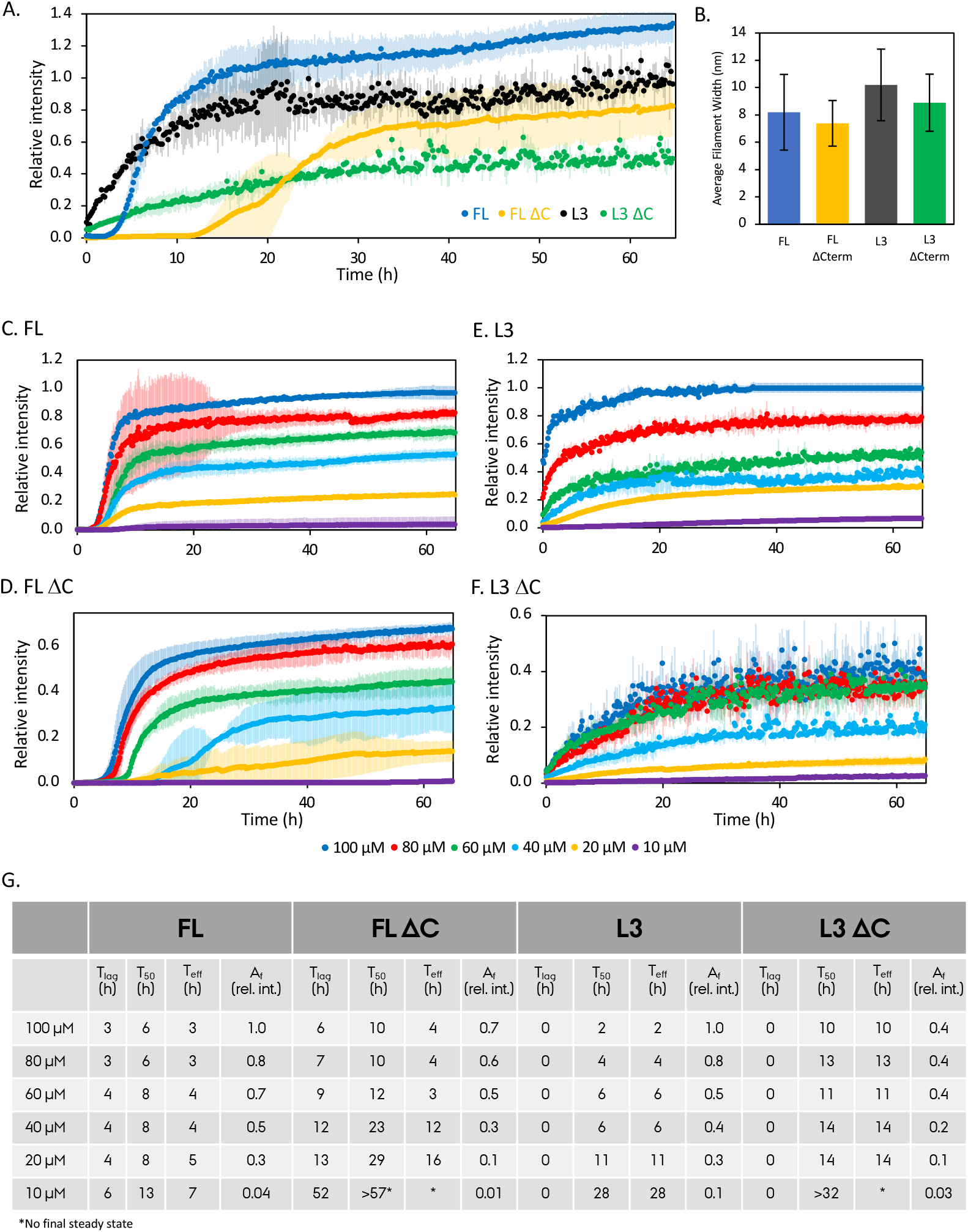
Concentration-dependent ThT assay reveal fibrillation kinetics being shaped by hierarchical layering and C-terminal modulation. Fibrillation kinetics of the FapC constructs (FL, FL ΔC, L3, L3 ΔC) were compared across protein concentrations. (A) ThT profiles at 40 μM, where the ThT fluorescence was normalized relative to L3. (B) The average fibril widths for FapC constructs. (C-F) Kinetics over a 10–100 μM range for each construct. Final ThT intensity was normalized to 100 μM L3 sample in E. Data reveal that both layer complexity and the C-terminal region modulate fibrillation onset and efficiency. (G) The kinetics parameters for all the measurements and samples. ThT samples that did not reach saturation were labeled with an asterisk and values are all approximations.

Concentration-dependent ThT kinetics revealed further distinctions, **Fig. 6B-F, SI3**. FL fibrillated robustly across concentrations from 100 to 10 μM, with T_lag_: 3-6 h and T_eff_: 3-7 h, **Fig. 6B,G**. In contrast, FL ΔC displayed a strong concentration dependence: T_lag_ and T_50_ progressively increased with decreasing concentration, with a sharp jump at 40 μM increasing T_lag_ from 9 to 12 h and quadrupling T_eff_ from 3 to 12 h, **Fig. 6C, G**. To the point where at 10 μM steady-state was not reached within the assay window of ∼65 h, **Fig. 6C, G**. L3 showed rapid fibrillation, with T_50_ decreasing from 28 h at 10 μM to just 2 h at 100 μM, **Fig. 6D, G**. No lag phase was observed across concentrations, except for a subtle delay at 10 μM, **Fig. SI3C**. End-point ThT signals remained similar to FL. L3 ΔC displayed intermediate kinetics: T_50_ values between 10-32 h from 100 to 10 μM, with no detectable lag phase but a plateau in final ThT values at 60 μM, **Fig. 6E**.

Together, these results demonstrate that the C-terminal segment and triple-layered architecture contribute independently to fibrillation. The C-terminus enhances elongation and yield across concentrations, while hierarchical assembly increases the nucleation barrier and introduces concentration-sensitive control via the lag phase. Our data suggests that FapC FL has been evolutionarily tuned to fibrillate most effectively, efficiently and consistently through the wide concentration range.

### High-resolution solution NMR quantifies the effect of C-terminus

To understand how the C-terminal region influences the conformational preferences of monomeric FapC, we used high-resolution solution NMR to determine chemical shift perturbations (CSPs) caused by C-terminal deletion throughout the FapC sequence, **Fig. 7**. All samples were uniformly ^15^N-labeled, and 2D ^1^H-^15^N HSQC spectra were acquired at 274 K and 600 MHz. Peak assignments were adapted from our previous NMR studies of FL and the L2R3C construct (residues 157-250), obtained under conditions but with 30 mM DTT.^49^ Over 90% of backbone amide peaks overlapped between FL and L3, and unassigned residues were excluded from further analysis, **Fig. 7C,D,SI5**.

**Figure 7.**
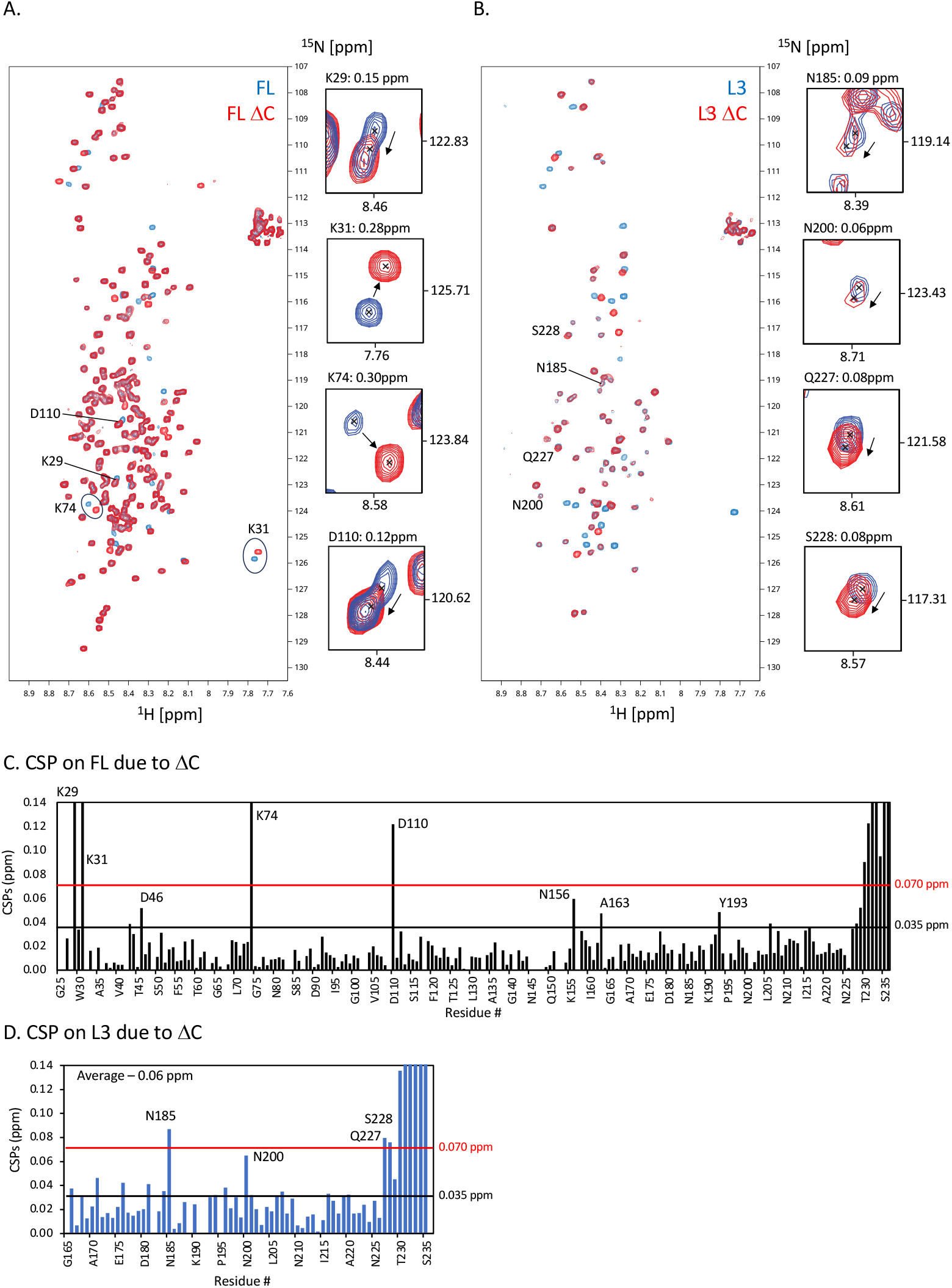
The effects of C-terminus deletion on FL and L3 FapC by solution-state NMR. ^1^H-^15^N HSQC spectrum analysis of FL (25-250), FL ΔC (25-236), L3 (165-250) and L3 ΔC (25-236). All experiments were measured at 274 K in 20 mM sodium phosphate, pH 7.8, 1 mM DSS, 10% D_2_O, and 0.02% sodium azide. (A) Overlay of FL (Blue) and FL ΔC (Red) spectra at 40 μM. (B) Overlay of L3 (blue) and L3 ΔC (red) spectra at 50 μM. (C, D) Composite CSP plots for ΔC vs. full-length constructs calculated by Δδ = ((5*Δδ^1H^)^2^ + (Δδ^15N^)^2^)^1/2^. CSP for FL ΔC compared to FL is shown in black, and L3 ΔC compared to L3 is shown in blue.

HSQC spectra of FL and L3 were overlayed with their respective ΔC-terminal variants, **Fig. 7A,B** to identify C-terminal-dependent perturbations. CSPs were calculated using Δδ = ((5*Δδ^1H^)^2^ + (Δδ^15N^)^2^)^1/2^ and categorized as: no CSP (<0.035 ppm), moderate CSP (0.035–0.07 ppm), or strong CSP (>0.07 ppm), **Fig. 7C,D**. C-terminal deletion led to widespread CSPs distributed across the FapC sequence, not limited to the truncation C-terminus region. In FL vs. FL ΔC, the largest perturbation was observed at K74 (0.3 ppm). Moderate-to-strong shifts were also seen at five charged residues (K29, K31, D46, D110, N156, N206), three hydrophobic residues (A43, A163, Y193), and one glycine (G216). In L3 vs. L3 ΔC, perturbations were seen at six charged residues (Q176, N181, N185, K190, Q196, N200) and one hydrophobic residue (L184), with the largest shift at N185 (0.09 ppm).

These CSPs shows that deletion of the C-terminal region perturbs residues far from the truncation site, consistent with long-range intramolecular interactions in the disordered monomeric state. Many of the perturbed residues are charged or polar, suggesting that electrostatic interactions between the disordered C-terminus and the core protein scaffold may guide early structural organization. These effects likely cooperate with previously described local secondary structure preferences to prime initial folding steps.^27,35^ Candidate residues in the C-terminus that may mediate these effects include the positively charged K247, disulfide-forming cysteines, and three polar residues (T238, T244, S246). Together, these findings support a model where the disordered C-terminal region stabilizes early folding intermediates, promoting productive fibril assembly even prior to nucleation.

## 4. Discussion

Functional amyloids must balance two competing demands: forming stable, β-sheet–rich fibrils while maintaining tightly regulated assembly. Unregulated aggregation can be toxic even in functional systems, and many known functional amyloids employ strict temporal and spatial control over their formation.^15-24^ Our study shows that this regulatory balance is encoded within the modular architecture and short C-terminal segment of the functional amyloid subunit FapC in *Pseudomonas*. Specifically, the single-layer L3 module fibrillates rapidly, while the FL construct exhibits a nucleation lag and strong concentration dependence, suggesting intrinsic checkpoints that modulate timing. L3 emerges as a highly efficient amyloidogenic unit that fibrillates with no lag phase, indicating a low nucleation barrier. This minimal construct behaves similarly to shorter pathogenic aβ fragments (aβ16–22) that aggregate faster due to reduced flexibility or lack of termini.^50^ In contrast, FL FapC exhibits a pronounced lag phase and sensitivity to coexisting species, highlighting how higher-order architecture imposes kinetic checkpoints to prevent premature aggregation.

The C-terminal region represents a second, independent regulator. Although not essential for fibrillation, its deletion slows nucleation, reduces fibrillation rate, and lowers yield, especially in monomer-only conditions. Solution NMR revealed long-range CSPs upon deletion, indicating that the C-terminal stabilizes monomers and facilitates early structural organization. This contrasts with pathologic amyloids like αSyn,^51^ where C-terminal deletion accelerates aggregation. Our data further show that oligomeric species modulate FapC assembly in architecture- and C-terminal-dependent ways. In FL, HMW oligomers promote fibrillation, likely through heterogeneous nucleation, while LMW species inhibit it by stabilizing off-pathway states. Modulation of nucleation and elongation by CsgC chaperone has been observed in functional amyloid CsgA from *E. coli*,^16,52^ and despite being different in nature may suggest that regulated assembly depends on selective progression and a possible control of intermediates. The effect is less pronounced in L3, underscoring that hierarchical structure and the presence of physical/chemical dimers in FL modulate this behavior. Notably, in ΔC constructs lacking these dimers, fibrillation relies primarily on monomer dynamics, distinguishing functional from pathologic amyloids as aβ and αSyn,^53-55^ which lack cysteines and often form non-productive off-pathway oligomeric intermediates not contributing to fibrillation.

Hierarchical layering and C-terminal regulation work synergistically but independently: the architecture imposes nucleation control, while the C-terminal region promotes elongation and yield. Accessory Fap proteins (FapA, FapB, and others) yet likely introduce an additional layer of regulation *in vivo*.^15^ Comparison across constructs reveals non-redundant contributions. L3 ΔC shows impaired fibrillation, confirming the intrinsic role of the C-terminal region even in minimal systems. Simplifying FL to L3 accelerates assembly but doesn’t restore efficiency if the C-terminus is absent. FL ΔC also shows reduced yield and prolonged lag, confirming the global regulatory function of the C-terminal segment.

A mechanistic model emerges, where FapC fibrillation initiates via the L3 segment forming a nucleation-competent core, with L2 and L1 completing the full three-layer fibril, as we showed previously by MD simulations.^27^ The C-terminal segment stabilizes folding intermediates and promotes elongation. HMW oligomers accelerate nucleation in FL, while LMW species divert monomers into off-pathway states. This model explains both the rapid assembly of L3 and the more regulated behavior of FL.

These principles distinguish functional from pathological amyloids. While both follow nucleation-elongation kinetics, β-solenoid functional amyloids incorporate modular, sequence-encoded checkpoints that ensure productive assembly. Pathological amyloids lack this regulation, often leading to toxic aggregates. FapC shows how the amyloid fold can be harnessed for biological function, not by avoiding aggregation, but by precisely controlling its timing and extent. In summary, our findings offer mechanistic insights that may guide targeted disruption of biofilm-associated amyloids. Rather than dissolving mature fibrils, interventions could inhibit L3 engagement, disrupt C-terminal-mediated elongation, or stabilize off-pathway species. Even modest disruptions may weaken biofilm integrity. Ultimately, regulation, not structure alone, may offer a promising path to modulating functional amyloids without disturbing proteostasis.

## Data Availability

All data can be requested from the corresponding author.

## Author Contribution

Conceptualization: UA; Methodology: CHB, AT, MU, XT, UA; Formal analysis and investigation: CHB, AT, MU, XT, UA; Writing - original draft preparation: CHB, UA; Funding acquisition: UA; Resources: UA; Supervision: UA. All authors reviewed the manuscript.

## Acknowledgements

The authors acknowledge support from the Department of Structural Biology at University of Pittsburgh School of Medicine (UPSOM) for access to the high-field NMR and EM facilities. Start-up funds by the UPSOM and a Competitive Medical Research Fund (CMRF) grant by the UPMC Health System to UA is acknowledged.

## Supplementary Figures

**Figure SI1.**
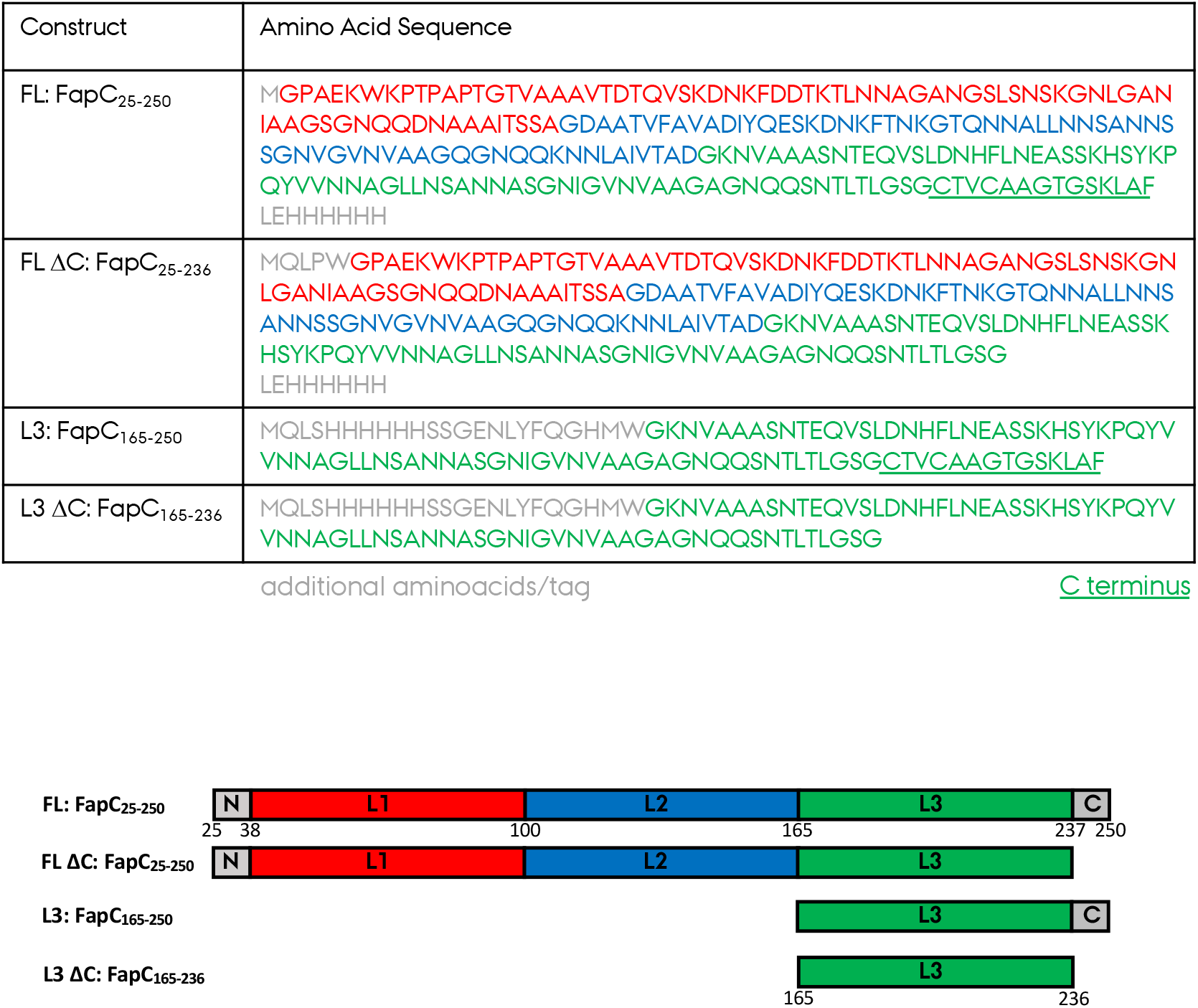
Full sequences of four FapC constructs used in this study.

**Figure SI2:**
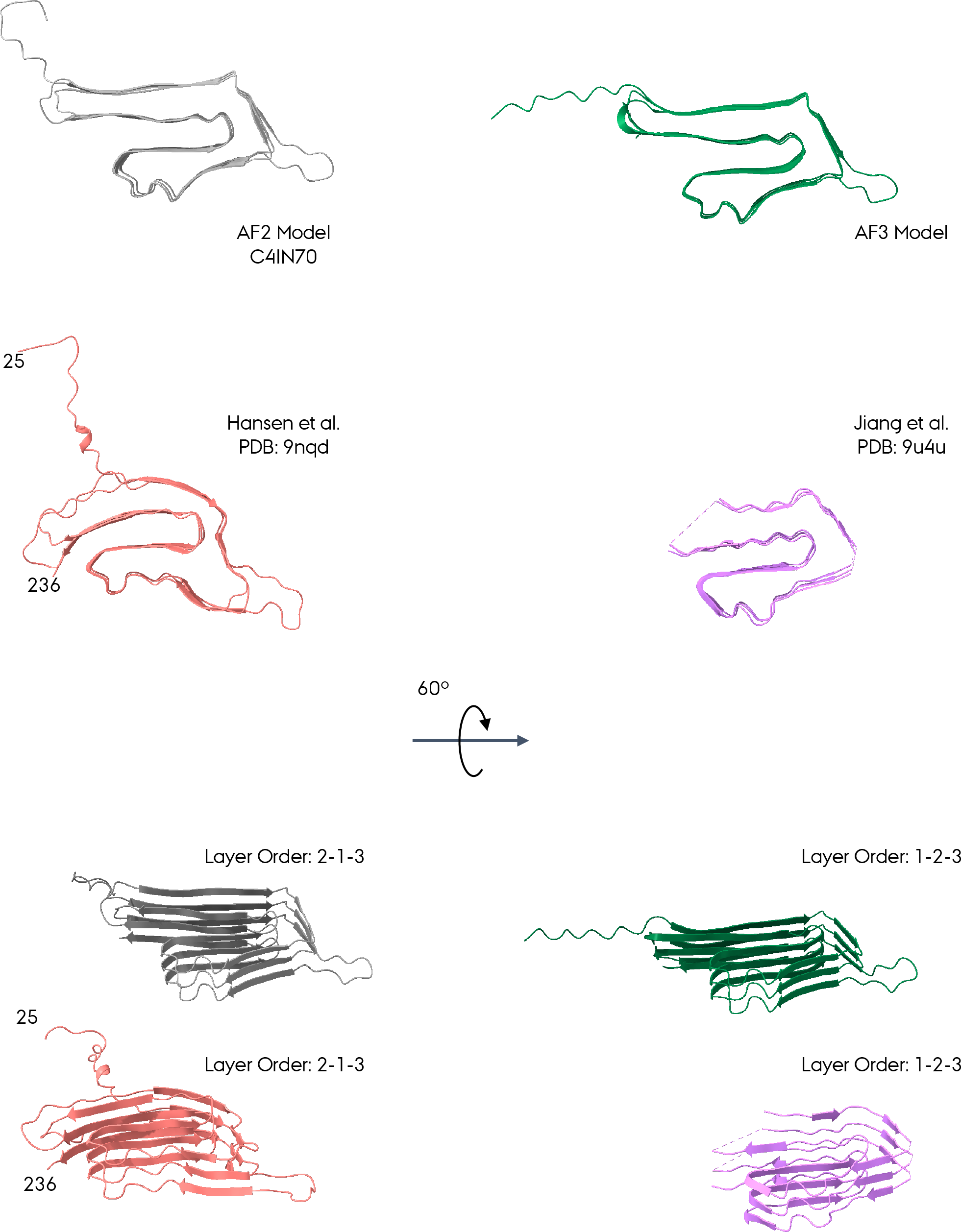
Experimentally determined (Hansen et al. PDB: 9nqd & Jiang et al. 9u4u) FapC_25-236_ structures and AlphaFold predicted FapC_25-236_ models (AF2: AF-C4IN70-F1 & AF3: AF-P0DXF5-F1).

**Figure SI3.**
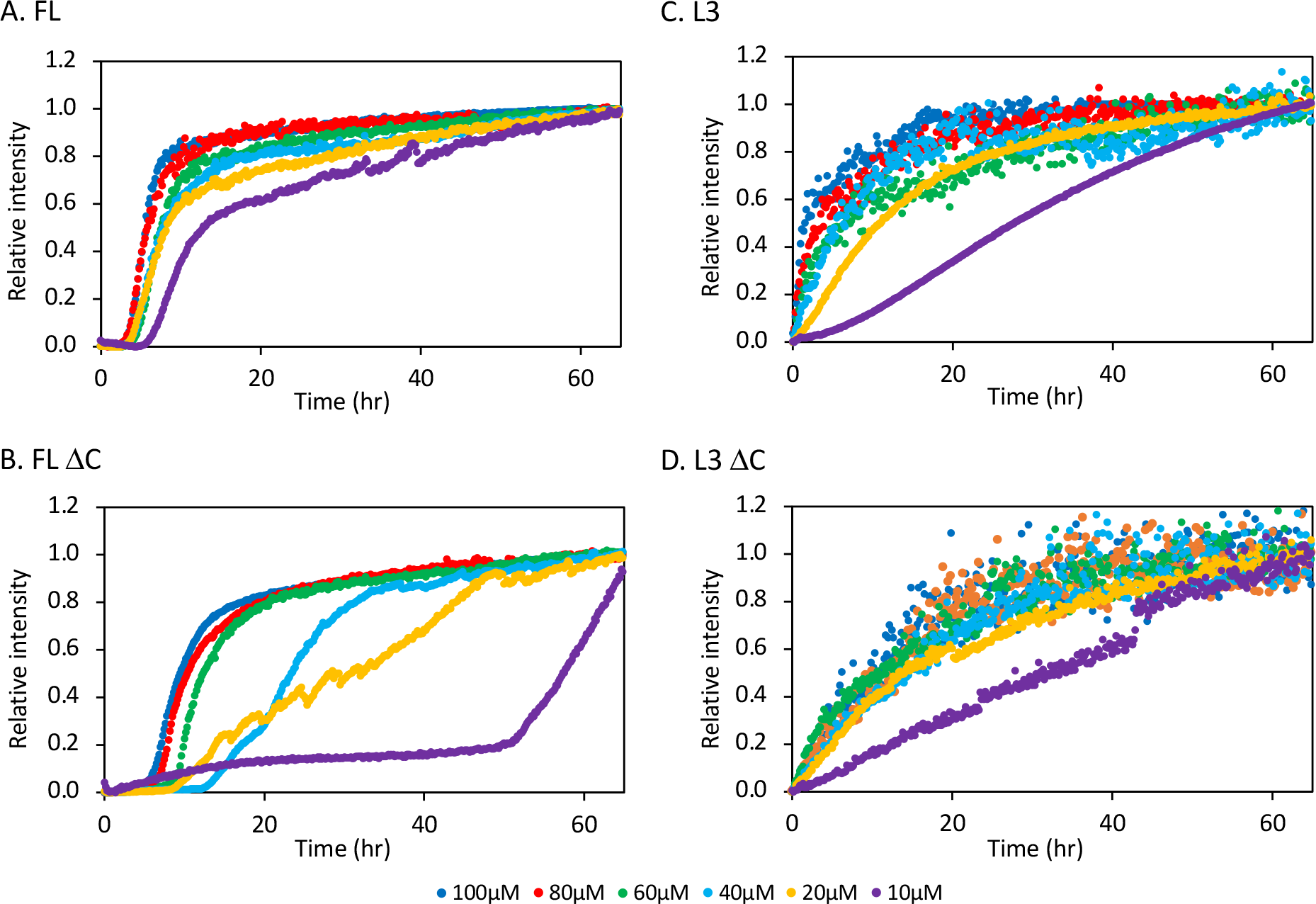
Absolute normalized ThT fluorescence for all four FapC constructs. The data is shown in the manuscript Figure 6 as absolute intensities.

**Figure SI4.**
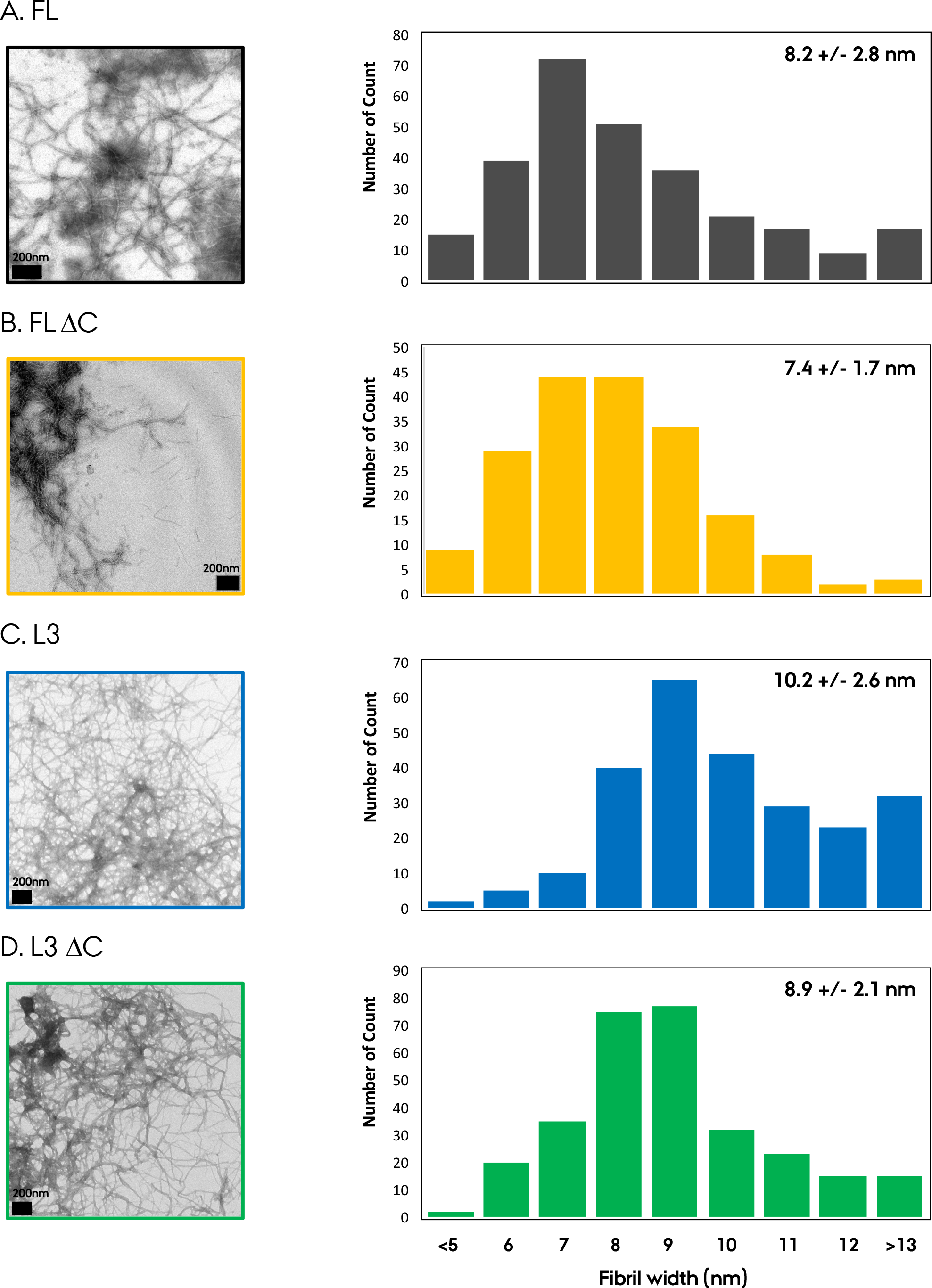
EM width measurements of amyloid fibrils of four different FapC samples.

**Figure SI5.**
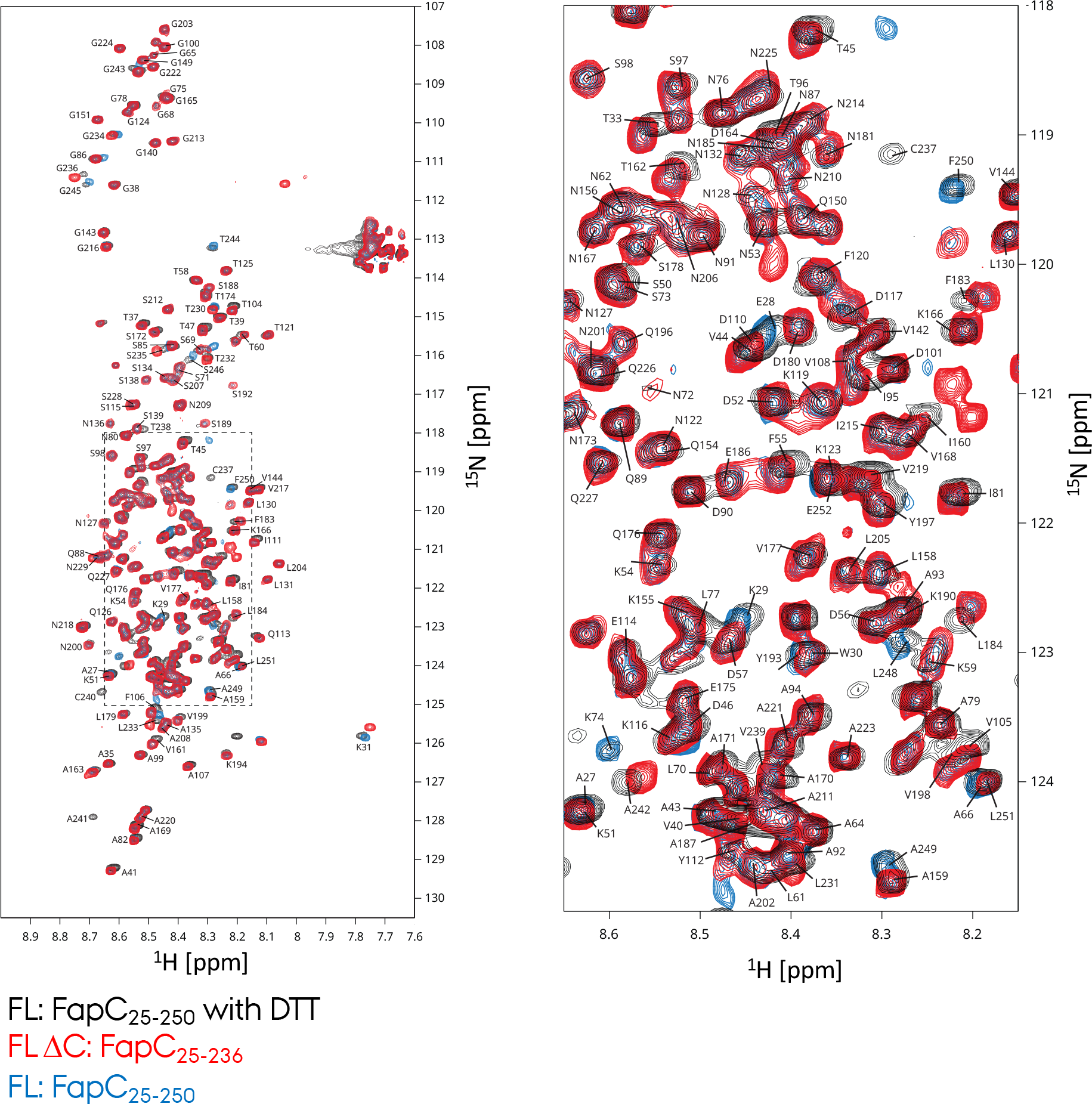
Solution-state NMR assignment comparison of the FL and FL ΔC FapC samples measured without DTT. These NMR spectra are compared to the FL FapC measured with DTT along with the assignments we reported previously (BMRB: #51793).

